# Bat Detective - Deep Learning Tools for Bat Acoustic Signal Detection

**DOI:** 10.1101/156869

**Authors:** Oisin Mac Aodha, Rory Gibb, Kate E. Barlow, Ella Browning, Michael Firman, Robin Freeman, Briana Harder, Libby Kinsey, Gary R. Mead, Stuart E. Newson, Ivan Pandourski, Stuart Parsons, Jon Russ, Abigel Szodoray-Paradi, Farkas Szodoray-Paradi, Elena Tilova, Mark Girolami, Gabriel Brostow, Kate E. Jones

## Abstract

1. Passive acoustic sensing has emerged as a powerful tool for quantifying anthropogenic impacts on biodiversity, especially for echolocating bat species. To better assess bat population trends there is a critical need for accurate, reliable, and open source tools that allow the detection and classification of bat calls in large collections of audio recordings. The majority of existing tools are commercial or have focused on the species classification task, neglecting the important problem of first localizing echolocation calls in audio which is particularly problematic in noisy recordings.
2. We developed a convolutional neural network (CNN) based open-source pipeline for detecting ultrasonic, full-spectrum, search-phase calls produced by echolocating bats (BatDetect). Our deep learning algorithms (CNN _FULL_ and CNN _FAST_) were trained on full-spectrum ultrasonic audio collected along road-transects across Romania and Bulgaria by citizen scientists as part of the iBats programme and labelled by users of <u>www.batdetective.org</u>. We compared the performance of our system to other algorithms and commercial systems on expert verified test datasets recorded from different sensors and countries. As an example application, we ran our detection pipeline on iBats monitoring data collected over five years from Jersey (UK), and compared results to a widely-used commercial system.
3. Here, we show that both CNN_FULL_ and CNN_FAST_ deep learning algorithms have a higher detection performance (average precision, and recall) of search-phase echolocation calls with our test sets, when compared to other existing algorithms and commercial systems tested. Precision scores for commercial systems were reasonably good across all test datasets (>0.7), but this was at the expense of recall rates. In particular, our deep learning approaches were better at detecting calls in road-transect data, which contained more noisy recordings. Our comparison of CNN_FULL_ and CNN_FAST_ algorithms was favourable, although CNN_FAST_ had a slightly poorer performance, displaying a trade-off between speed and accuracy. Our example monitoring application demonstrated that our open-source, fully automatic, BatDetect CNN_FAST_ pipeline does as well or better compared to a commercial system with manual verification previously used to analyse monitoring data.
4. We show that it is possible to both accurately and automatically detect bat search-phase echolocation calls, particularly from noisy audio recordings. Our detection pipeline enables the automatic detection and monitoring of bat populations, and further facilitates their use as indicator species on a large scale, particularly when combined with automatic species identification. We release our system and datasets to encourage future progress and transparency.

## Introduction

There is a critical need for robust and accurate tools to scale up biodiversity monitoring and to manage the impact of anthropogenic change (Cardinale *et al.* 2012; Turner 2014). Modern hardware for passive biodiversity sensing such as camera trapping and audio recording now enables the collection of vast quantities of data relatively inexpensively. In recent years, passive acoustic sensing has emerged as a powerful tool for understanding trends in biodiversity (Sueur *et al.* 2009; Blumstein *et al.* 2011; Marques *et al.* 2013; Penone *et al.* 2013). Monitoring of bat species and their population dynamics can act as an important indicator of ecosystem health as they are particularly sensitive to habitat conversion and climate change (Jones *et al.* 2013). Close to 80% of bat species emit ultrasonic pulses, or echolocation calls, to search for prey, avoid obstacles, and to communicate (Schnitzler, Moss & Denzinger 2003). Acoustic monitoring offers a passive, non-invasive, way to collect data about echolocating bat population dynamics and the occurrence of species, and it is increasingly being used to survey and monitor bat populations (Jones *et al.* 2013; Barlow *et al.* 2015; Newson, Evans & Gillings 2015).

Despite the obvious advantages of passive acoustics for monitoring echolocating bat populations, its widespread use has been hampered by the challenges of robust identification of acoustic signals, generation of meaningful statistical population trends from acoustic activity, and engaging a wide audience to take part in monitoring programmes (Walters *et al.* 2013). Recent developments in statistical methodologies for estimating abundance from acoustic activity (Marques *et al.* 2013; Lucas *et al.* 2015; Stevenson *et al.* 2015), and the growth of citizen science networks for bats (Barlow *et al.* 2015; Newson, Evans & Gillings 2015) mean that efficient and robust audio signal processing tools are now a key priority. However, tool development is hampered by a lack of large scale species reference audio datasets, intraspecific variability of bat echolocation signals, and radically different recording devices being used to collect data (Walters *et al.* 2013).

To date, most full-spectrum acoustic identification tools for bats have focused on the problem of species classification from search-phase echolocation calls (Walters *et* al. 2013). Existing methods typically extract a set of audio features (such as call duration, mean frequency, and mean amplitude) from high quality search-phase echolocation call reference libraries to train machine learning algorithms to classify unknown calls to species (Parsons & Jones 2000; Russo & Jones 2002; Skowronski & Harris 2006; Armitage & Ober 2010; Walters *et al.* 2012; Walters *et al.* 2013; Zamora-Gutierrez *et al.* 2016). Instead of using manually defined features, another set of approaches attempt to learn representation directly from spectrograms (Stowell & Plumbley 2014; Stathopoulos *et al.* 2017). Localising audio events in time (defined here as ‘detection’), is an important challenge in itself, and is often a necessary pre-processing step for species classification (Stowell *et al.* 2016). Additionally, understanding how calls are detected is critical to quantifying any biases which may impact estimates of species abundance or occupancy (Clement *et al.* 2014b; Lucas *et* al. 2015). For example, high levels of background noise, often found in highly disturbed anthropogenic habitats such as cities, may have a significant impact on the ability to detect signals in recordings and lead to a bias in population estimates.

Detecting search-phase calls by manual inspection of spectrograms tends to be subjective, highly dependent on individual experience, and its uncertainties are difficult to quantify (Skowronski & Fenton 2008). There are a number of automatic detection tools now available which use a variety of methods, including amplitude threshold filtering, locating areas of smooth frequency change, detection of set search criteria, or based on a cross-correlation of signal spectrograms with a reference spectrogram (see review in Walters *et al.* 2013). While there are some studies that analyse the biases of automated detection (and classification) tools (Jennings, Parsons & Pocock 2008; Adams *et al.* 2012; Clement *et al.* 2014a; Fritsch & Bruckner 2014; Russo & Voigt 2016; Rydell *et al.* 2017), this is generally poorly quantified, and in particular, there is very little published data available on the accuracy of many existing closed source commercial systems. Despite this, commercial systems are commonly used in bat acoustic survey and monitoring studies, albeit often with additional manual inspection (Barlow *et al.* 2015; Newson, Evans & Gillings 2015). This reliance on poorly documented algorithms is scientifically undesirable, and manual detection of signals is clearly not scalable for national or regional survey and monitoring. In addition, there is the danger that manual detection and classification introduces a bias towards the less noisy and therefore more easily identifiable calls. To address these limitations, a freely available, transparent, fast, and accurate detection algorithm that can also be used alongside other classification algorithms is highly desirable.

Here, we develop an open source system for automatic bat search-phase echolocation call detection (i.e. localisation in time) in noisy, real world, recordings. We use the latest developments in machine learning to directly learn features from the input audio data using supervised deep convolutional neural networks (CNNs) (LeCun *et al.* 1998). CNNs have been shown to be very successful for classification and detection of objects in images (Krizhevsky, Sutskever & Hinton 2012; Girshick *et al.* 2014). They have also been applied to various audio classification tasks (Piczak 2015; Hershey *et al.* 2016; Salamon & Bello 2016), along with human speech recognition (Hinton *et al.* 2012; Hannun *et al.* 2014). Although CNNs are now starting to be used for bioacoustic signal detection and classification tasks in theoretical or small-scale contexts (e.g. bird call detection) (Goeau *et al.* 2016), to date there have been no application of CNN-based tools for bat monitoring. This is mainly due to a lack of sufficiently large labelled bat audio datasets for use as training data. To overcome this, we use data collected and annotated by thousands of citizen scientists as part of our Indicator Bats Programme (Jones *et al.* 2013) and Bat Detective (<u>www.batdetective.org</u>). We validate our system on three different challenging test datasets from Europe which represent realistic use cases for bat surveys and monitoring programmes, and we present an example real-world application of our system on five years of monitoring data collected in Jersey (UK).

## Materials and Methods

### ACOUSTIC DETECTION PIPELINE

We created a detection system to determine the temporal location of any search-phase bat echolocation calls present in ultrasonic audio recordings. Our detection pipeline consisted of four main steps (Figure 1) as follows: (1) *Fast Fourier Transform Analysis* - Raw audio (Figure 1a) was converted into a log magnitude spectrogram (FFT window size 2.3 milliseconds, overlap of 75%, with Hanning window), retaining the frequency bands between 5kHz and 135kHz (Figure 1b). Recordings with a sampling rate of 44.1kHz, time expansion factor of 10, and 2.3ms FFT window, resulted in a window size of 1,024 samples. We used spectrograms rather than raw audio for analysis, as it provides an efficient means of dealing with audio that has been recorded at different sampling rates. Provided the frequency and time bins of the spectrogram are of the same resolution, audio with different sampling rates can be input into the same network. (2) *De-noising* – We used the de-noising method of (Aide *et al.* 2013) to filter out background noise by removing the mean amplitude in each frequency band (Figure 1c), as this significantly improved performance. (3) *Convolutional Neural Network Detection*– We created a convolutional neural network (CNN) that poses search-phase bat echolocation call detection as a binary classification problem. Our CNN_FULL_ consisted of three convolution and max pooling layers, followed by one fully connected layer (see Supplementary Information Methods for further details). We halved the size of the input spectrogram to reduce the input dimensionality to the CNN which resulted in an input array of size of 130 frequency bins by 20 time steps, corresponding to a, fixed length, detection window size of 23ms. We applied the CNN in a sliding window fashion, to predict the presence of a search-phase bat call at every instance of time in the spectrogram (Figure 1d). As passive acoustic monitoring can generate large quantities of data, we required a detection algorithm that would run faster than real time. While CNNs produce state of the art results for many tasks, naïve application of them for detection problems at test time can be extremely computationally inefficient (Girshick *et al.* 2014). So, to increase the speed of our system we also created a second, smaller CNN which included fewer model weights that can be run in a fully convolutional manner (CNN_FAST_) (Supplementary Information Methods, Supplementary Information Figure S1). (4) *Call Detection Probabilities* – The probabilistic predictions produced by the sliding window detector tended to be overly smooth in time (Figure 1d). To localise the calls precisely, we converted the probabilistic predictions into individual detections using a non-maximum suppression to return the local maximum for each peak in the output prediction (Figure 1e). These local maxima corresponded to the predicted locations of the start of each search-phase bat echolocation call, with associated probabilities, and were exported as text files.

**Figure 1.**
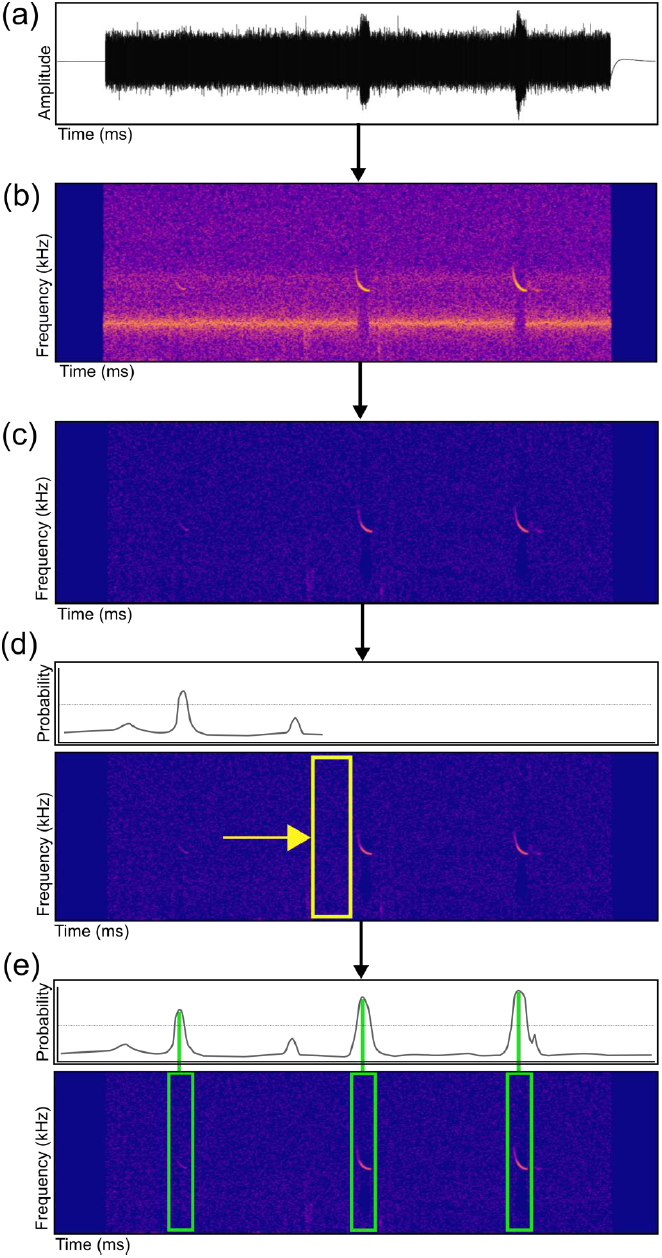
Detection pipeline for search-phase bat echolocation calls. (a) Raw audio files are converted into a spectrogram using a Fast Fourier Transform (b). Files are de-noised (c), and a sliding window Convolutional Neural Network (CNN) classifier (d, yellow box) produces a probability for each time step. Individual call detection probabilities using non-maximum suppression are produced (e, green boxes), and the time in file of each prediction along with the classifier probability are exported as text files.

### ACOUSTIC TRAINING DATASETS

We trained our BatDetect CNNs using a subset of full-spectrum time-expanded (TE) ultrasonic acoustic data recorded between 2005-2011 along road-transects by citizen scientists as part of the Indicator Bats Programme (iBats) (Jones *et al.* 2013) (see Supplementary Information Methods for detailed data collection protocols). During surveys, acoustic devices (Tranquility Transect, Courtplan Design Ltd, UK) were set to record using a TE factor of 10, a sampling time of 320ms, and sensitivity set on maximum, giving a continuous sequence of ‘snapshots’, consisting of 320ms of silence (sensor listening) and 3.2s of TE audio (sensor playing back x 10). As sensitivity was set at maximum, and no minimum amplitude trigger mechanism was used on the recording devices, our recorded audio data contained many instances of low amplitude and faint bat calls, as well as other night-time ‘background’ noises such as other biotic, abiotic, and anthropogenic sounds.

We generated annotations of the start time of search-phase bat echolocation calls in the acoustic recordings by uploading the acoustic data to the Zooniverse citizen science platform (<u>www.zooniverse.org</u>) as part of the Bat Detective project (<u>www.batdetective.org</u>), to enable public users to view and annotate them. The audio data were first split up into 3.84s long sound clips to include the 3.2s of TE audio and buffered by sensor-listening silence on either side. We then uploaded each sound clip as both a wav file and a magnitude spectrogram image (represented as a 512×720 resolution image) onto the Bat Detective project website. As the original recordings were time-expanded, therefore reducing the frequency, sounds in the files were in the audible spectrum and could be easily heard by users. Users were presented with a spectrogram and its corresponding audio file, and asked to annotate the presence of bat calls in each 3.84s clip (corresponding to 320ms of real-time recordings) (Supplementary Information Figure S2). After an initial tutorial (Supplementary Information Video 1), users were instructed to draw bounding boxes around the locations of bat calls within call sequences and to annotate them as being either: (1) search-phase echolocation calls; (2) terminal feeding buzzes; or (3) social calls. Users were also encouraged to annotate the presence of insect vocalisations and non-biotic mechanical noises.

Between Oct 2012 and Sept 2016, 2,786 users (including only the number of users which had registered with the site and did more than five annotations) listened to 127,451 unique clips and made 605,907 annotations. 14,339 of these clips were labelled as containing a bat call, with 10,272 identified as containing search-phase echolocation calls. Due to the inherent difficultly of identifying bat calls and the inexperience of some of our users, we observed a large number of errors in the annotations provided. To overcome this problem, we visually inspected a subset of the annotations from our most active users and found that they produced high quality annotations. As a result, we choose annotations from the top user who had viewed 46,508 unique sound clips and had labelled 3,364 clips as containing bat search-phase echolocation calls (a representative sample is shown in Supplementary Information Figure S3). From this we randomly selected a training set of 2,812 clips, consisting of 4,782 individual search-phase echolocation call annotations from Romania and Bulgaria, with which to train the CNNs (corresponding to data from 347 road-transect sampling events of 137 different transects collected between 2006 and 2011) (Figure 2a). Data were chosen from these countries as they contain the majority of the most commonly occurring bat species in Europe (IUCN 2017). This training set was used for all experiments. The remaining annotated clips from the same user were used to create one of our test sets, iBats Romania and Bulgaria (Figure 2a and see below). Occasionally, call harmonics and the associated main call were sometimes labelled with different start times in the same audio clip. To address this problem, we automatically merged annotations that occurred within 6 milliseconds of each other, making the assumption that they belonged to the same call.

**Figure 2.**
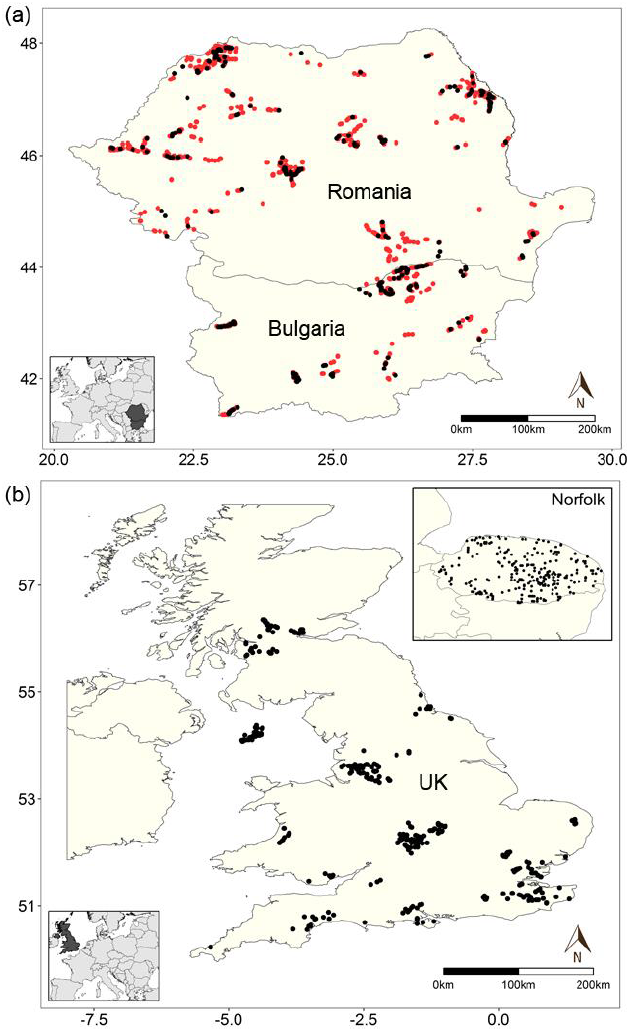
Spatial distribution of the BatDetect CNNs training and testing datasets. (a) Location of training data for all experiments and one test dataset in Romania and Bulgaria (2006-2011) from time-expanded (TE) data recorded along road transects by the Indicator Bats Programme (iBats) (Jones *et al.* 2013), where red and black points represent training and test data, respectively. (b) Locations of additional test datasets from TE data recorded as part of iBats car transects in the UK (2005-2011), and from real-time recordings from static recorders from the Norfolk Bat Survey from 2015 (inset). Points represent the start location of each snapshot recording for each iBats transect or locations of static detectors for the Norfolk Bat Survey.

### ACOUSTIC TESTING DATASETS AND EVALUATION

To evaluate the performance of the detection algorithms, we created three different test datasets of approximately the same size (number and length of clips) (Figure 2a-b, Supplementary Information Table S1). These datasets were chosen to represent three different realistic use cases commonly used for bat surveys and monitoring programmes and included data collected both along road-transects (resulting in noisier audio), and using static ultrasonic detectors. The test sets were as follows: (1) *iBats Romania and Bulgaria* - audio recorded from the same region, by the same individuals, with the same equipment, and sampling protocols as the training set, corresponding to 161 sampling events of 81 different transect routes; (2) *iBats UK* - audio recorded from a different region (corresponding to data from 176 sampling events of 111 different transects recorded between 2005-2011 in the United Kingdom, chosen randomly), by different individuals, using the same equipment type, and identical sampling protocols as part of the iBats programme (Jones *et al.* 2013) as the training set; and (3) *Norfolk Bat Survey* - audio recorded from a different region (Norfolk, UK), by different individuals, using different equipment types (SM2BAT+ Song Meter, Wildlife Acoustics) and different protocols (static devices from random sampling locations) as part of the Norfolk Bat Survey (Newson, Evans & Gillings 2015) in 2015. These data corresponded to 381 sampling events from 246 static recording locations (1km^2^ grid cells), randomly chosen. The start times of the search-phase echolocation calls in these three test sets were manually extracted. For ambiguous calls, we consulted two experts, each with over 10 years of experience with bat acoustics.

As these data contained a significantly greater proportion of negative (non-bat calls) as compared to positive examples (bat calls), standard error metrics used for classification such as overall accuracy were not suitable for evaluating detection. Instead, we report the interpolated average precision and recall of each method displayed as a precision-recall curve (Everingham *et al.* 2010). Precision was calculated as the number of true positives divided by the sum of both true and false positives. We consider a detection to be a true positive if it occurred within 10ms of the expert annotated start time of the search-phase echolocation call. Recall was measured as the overall fraction of calls that were present in the audio that were correctly detected. Curves were obtained by thresholding the detection probabilities from zero to one and recording the precision and recall at each threshold. Algorithms that did not produce a continuous output were represented as a single point on the precision-recall curves. We also report recall at 0.95 precision, a metric that measures the fraction of calls that were detected while accepting a false positive rate of 5%. Thus a detection algorithm gets a score of zero if it was not capable of retrieving any calls with a precision greater than 0.95.

We compared the performance of both BatDetect CNNs to three existing closed-source commercial detection systems: (1) SonoBat (version 3.1.7p) (Szewczak 2010); (2) SCAN’R version 1.7.7. (Binary Acoustic Technology 2014); and (3) Kaleidoscope (version 4.2.0 alpha4) (Wildlife Acoustics 2012). For SonoBat, calls were extracted in batch mode. We set a maximum of 100 calls per file (there are never more than 20 search-phase calls in a test file), and set ‘acceptable call quality’ and ‘skip calls below this quality’ parameters both to zero, and used an auto filter of 5KHz. For SCAN‘R, we used standard settings as follows: setting minimum and maximum frequency cut off at 10 kHz and 125 kHz, respectively; minimum call duration at 0.5 ms; and minimum trigger level of 10 dB. We used Kaleidoscope in batch mode, setting ‘frequency range’ to 15-120kHz, ‘duration range’ to 0-500ms, ‘maximum inter-syllable’ to 0ms, and ‘minimum number of pulses’ to 0. We also compared two other detection algorithms that we implemented ourselves, which are representative of typical approaches used for detection in audio files and in other bat acoustic classification studies: (4) Segmentation-an amplitude thresholding segmentation method (Lasseck 2014), this is related to the approach of (Bas, Bas & Julien 2017); and (5) Random Forest – a random forest-based classifier (Breiman 2001). Where relevant, the algorithms for (4) and (5) used the same processing steps as the BatDetect CNNs. For the Segmentation method, we thresholded the amplitude of the input spectrogram resulting in a binary segmentation. Regions that were greater than the threshold *S*_*t*_, and bigger than size *S*_*r*_, were considered as positive instances. We chose the values of *S*_*t*_ and *S*_*r*_ on the iBats (Romania and Bulgaria) test dataset that gave the best test results to quantify its best case performance. For the *Random Forest* algorithm, we used the raw amplitude values from the gradient magnitude of the log magnitude spectrogram as features. We compared the total processing time for each of our own algorithms, and timings were calculated on a desktop with an Intel i7 processor, 32Gb of RAM, and a Nvidia GTX 1080 GPU. With the exception of the BatDetect CNN _FULL_, which used a GPU at test time, all the other algorithms were run on the CPU.

### ECOLOGICAL MONITORING APPLICATION

To demonstrate the performance of our method in a large-scale ecological monitoring application, we compared the number of bat search-phase echolocation calls found using our BatDetect CNN_FAST_ algorithm to those produced from a commonly used commercial package using SonoBat (version 3.1.7p) (Szewczak 2010) as a baseline, using monitoring data collected in iBats programme in Jersey, UK from 2011-2015. Audio data was collected twice yearly (July and August) from 11 road-transect routes of approximately 40km by volunteers using the iBats protocols (Supplementary Information, Supplementary Methods), corresponding to 5.7 days of continuous TE audio over five years (or 13.75 hours of real-time data). For the BatDetect CNN_FAST_ analysis, we ran the pipeline as described above, using a conservative probabilistic threshold of 0.90 (so as to only include high precision predictions). Computational analysis timings for the CNN_FAST_ for this dataset were calculated as before. For the comparison to SonoBat, we used the results from an existing real-world analysis in a recent monitoring report (Walters, Browning & Jones 2016), where the audio files were first split into 1 min recordings, and then SonoBat was used to detect search-phase calls and to fit a frequency-time trend line to the shape of the call (Walters, Browning & Jones 2016). All fitted lines were visually inspected and calls where the fitted line included background noise or echoes, were rejected. Typically, monitoring analyses group individual calls into sequences (a bat pass) before analysis. To replicate that here in both analyses, individual calls were assumed to be part of the same call sequence (bat pass) if they occurred within the same 3.84s sound clip and if the sequence continued into subsequent sound clips. We compared number of bat calls and passes detected per transect sampling event across the two analyses methods using generalized linear mixed models (GLMM) using lme4 (Bates *et al.* 2015) in R v. 3.3.3 (R Development Core Team 2009) in order to control for the spatial and temporal non-independence of our survey data (Poisson GLMM including analysis method as a fixed effect and sampling event, transect route and date as random effects).

## Results

### ACOUSTIC DETECTION PERFORMANCE

Both versions of our BatDetect CNN algorithm outperformed all other algorithms and commercial systems tested, with consistently higher average precision scores and recall rates across the three different test datasets (Table 1, Figure 3a-c). In particular, the CNNs detected a substantially higher proportion of search-phase calls at 0.95 precision (maximum 5% false positives) (Table 1). All the other algorithms underestimated the number of search-phase echolocation calls in each dataset, except Segmentation, which produced high recall rates but with low precision (a high number of false positives). The CNNs relative improvement compared to other methods was higher on the road transect datasets (iBats Romania & Bulgaria; iBats UK; Table 1, Figure 3a-b). Overall the performance of CNN_FAST_ was slightly worse than the larger CNN_FAST_ across all test datasets, apart from a higher recall rates at 0.95 precision in the static Norfolk Bat Survey dataset (Figure 3c, Table 1). Precision scores for all commercial systems (SonoBat, SCAN’R and Kaleidoscope) were reasonably good across all test datasets (>0.7) (Figure 3a-c). However, this was at the expense of recall rates, which were consistently lower than for the CNNs and Random Forest, where the maximum recall rates were 44-60% of known calls detected (Figure 3c). The recall rates fell to a maximum of 25% of known calls for the road transect datasets (Figure 3a-b).

**Table 1.**
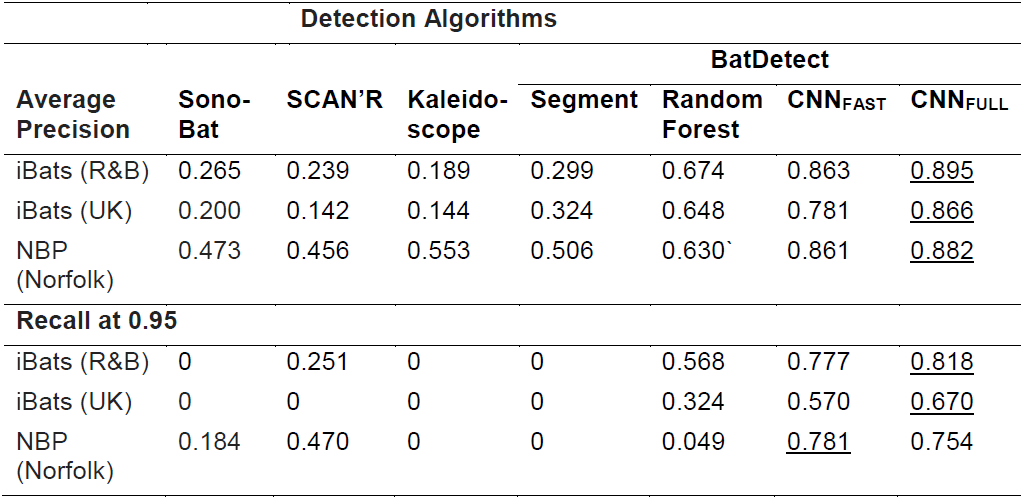
Average precision and recall results for bat search-phase call detection algorithms across three different test sets iBats Romania and Bulgaria; iBats UK; and Norfolk Bat Survey. Large numbers indicate better performance. Recall results are reported at 0.95 precision, where zero indicates that the detector algorithm was unable to achieve a precision greater than 0.95 at any recall level. The results for the best performing algorithm are underlined. Details of the test datasets and detection algorithms are given in the text.

**Figure 3.**
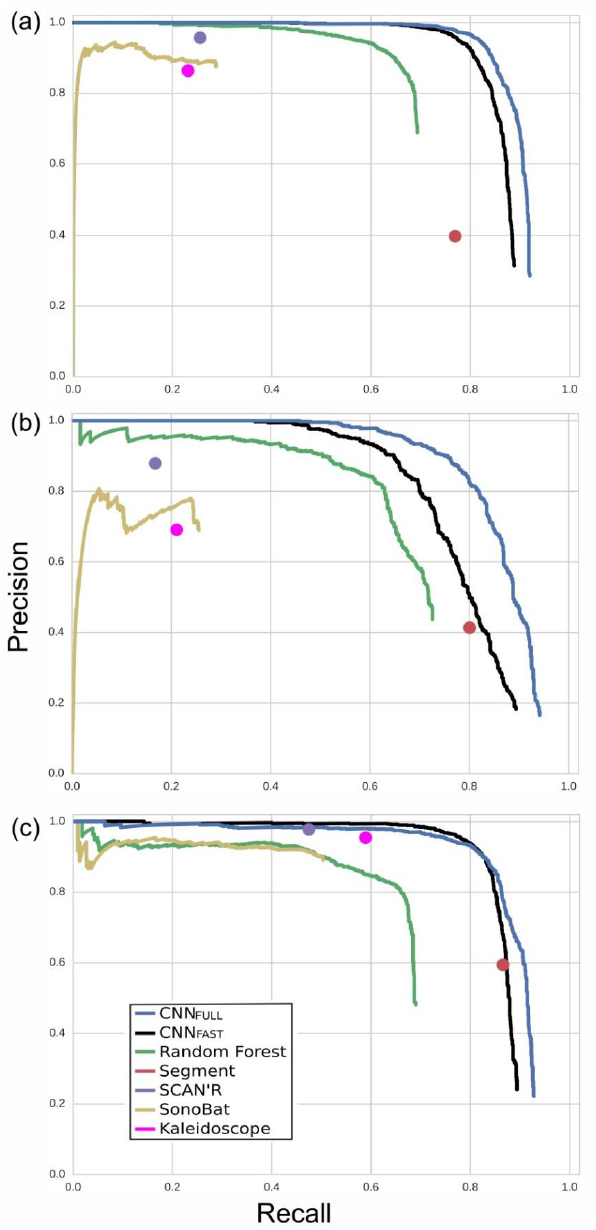
Precision-recall curves for bat search-phase call detection algorithms across three testing datasets; (a) iBats Romania and Bulgaria; (b) iBats UK; and (c) Norfolk Bat Survey. Curves were obtained by sweeping the output probability for a given detector algorithm and computing the precision and recall at each threshold. The commercial systems or algorithms that did not return a continuous output or probability (SCAN’R, Segment, and Kaleidoscope) were depicted as a single point.

CNN_FULL_, CNN_FAST_, Random Forest, and the Segmentation algorithms took 53, 9.5, 11, and 17 seconds respectively, to run the full detection pipeline on the 3.2 minutes of full spectrum iBats Romania and Bulgaria test dataset. Compared to CNN_FULL_ there was therefore a significant decrease in the time required to perform detection using CNN_FAST_, which was also the fastest of our methods overall. Notably, close to 50% of the CNN runtime was spent generating the spectrograms for detection, making this the most computationally expensive stage in the pipeline.

### ECOLOGICAL MONITORING APPLICATION

Our BatDetect CNN_FAST_ algorithm detected a significantly higher number of bat echolocation search-phase calls per transect sampling event, across 5 years of road transect data from iBats Jersey, compared to using SonoBat (CNN_FAST_ mean=107.69, sd=48.01; SonoBat mean=64.95, sd=28.53, Poisson GLMM including sampling event, transect route and date as random effects p<2e-^16^, n=216) (Figure 4, Supplementary Information Table S2). The differences between the two methods for bat passes was much smaller per sampling event, although CNN_FAST_ still detected significantly more passes per transect recording (CNN_FAST_ mean=29.57, sd=11.26; SonoBat mean=27.27, sd=10.85; Poisson GLMM including sampling event, transect route and date as random effects p=0.00143, n=216) (Figure 4, Supplementary Information Table S2). Running only on the CPU, the CNN_FAST_ algorithm took 24s to process each full transect of time-expanded audio (over 150 times real time).

**Figure 4.**
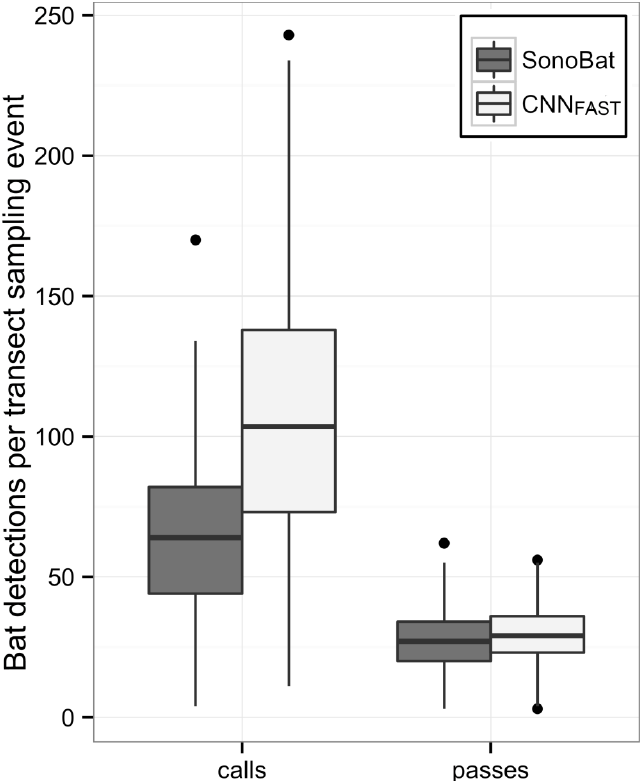
Comparison of the predicted bat detections (calls and passes) for two different acoustic systems using monitoring data collected from Jersey, UK. Acoustic systems used were SonoBat (version 3.1.7p) (Szewczak 2010) using analysis in (Walters, Browning & Jones 2016), and BatDetect CNNFAST using a probability threshold of 0.90. Detections are shown within each box plot, where the black line represents the mean across all transect sampling events from 2011-2015, boxes represent the middle 50% of the data, whiskers represent variability outside the upper and lower quartiles, with outliers plotted as individual points. See text for definition of a bat pass.

## Discussion

The BatDetect deep learning algorithms show a higher detection performance (average precision and recall) for search-phase echolocation calls with the test sets, when compared to other existing algorithms and commercial systems. In particular, our algorithms were better at detecting calls in road-transect data, which tend to contain more noisy recordings, suggesting that these are extremely useful tools for measuring bat abundance and occurrence in such datasets. Road-transect acoustic monitoring is a useful technique to assess bat populations over large areas and programmes have now been established by government and non-government agencies in many different countries (e.g., Roche *et al.* 2011; Jones *et al.* 2013; Whitby *et al.* 2014; Loeb *et al.* 2015; Azam *et al.* 2016). Noisy sound environments are also likely to be a problem for other acoustic bat monitoring programmes. For example, with the falling cost and wider availability of full-spectrum audio equipment, the range of environments being acoustically monitored is increasing, including noisy urban situations (Lintott *et al.* 2015; Merchant *et al.* 2015). Individual bats further from the microphone are less likely to be detected as their calls are fainter, and high ambient noise levels increase call masking and decrease call detectability. Additionally, a growth in open-source sensor equipment for bat acoustics using very cheap MEMs microphones (Whytock & Christie 2017) may also require algorithms able to detect bats in lower quality recordings, which may have a lower signal to noise ratio or a reduced call band-width due to frequency-dependent loss. Our open-source, well documented algorithms enable biases and errors to be directly incorporated into any acoustic analysis of bat populations and dynamics (e.g. occupancy models (e.g., Clement *et al.* 2014b). The detections with BatDetect can be directly used as input for population monitoring programmes when species identification is difficult such as the tropics, or to other CNN systems to determine bat species identity when sound libraries are available.

Our result that deep learning networks consistently outperformed other baselines, is consistent with the suggestion that CNNs offer substantially improved performance over other supervised learning methods for acoustic signal classification (Goeau *et al.* 2016). The major improvement of both CNNs over Random Forest and the three commercial systems was in terms of recall, i.e. increasing the proportion of detected bat calls in the test datasets. Although the precision of the commercial systems was often relatively high, the CNNs were able to detect much fainter and partially noise-masked bat calls that were missed by the other methods, with fewer false positives, and very quickly, particularly with CNN_FAST_. Previous applications of deep learning networks to bioacoustic and environmental sound recognition have used small and high-quality datasets (e.g., Goeau *et al.* 2016; Salamon & Bello 2016). However, our results show that, provided they are trained with suitably large and varied training data, deep learning networks have good potential for applied use in real-world heterogeneous datasets that are characteristic of acoustic wildlife monitoring (involving considerable variability in both environmental noise and distance of animal from sensor). Our comparison of CNN_FULL_ and CNN_FAST_ detectors was favourable, although CNN_FAST_ had a slightly poorer performance showing a trade-off between speed and accuracy. This suggests that CNN_FAST_ could potentially be adapted to work well with on-board low power devices (e.g. Intel’s Edison device) to deliver real-time detections. Avoiding the spectrogram generation stage entirely and using the raw audio samples as input (van den Oord *et al.* 2016), could also speed up performance of the system in the future, as currently over 50% of the CNN test time is taken up by computing spectrograms.

While our results have been validated on European bats, no species or region-specific knowledge, or particular acoustic sensor system is directly encoded into our system, making it possible to easily generalise to other systems (e.g. frequency division recordings), regions and species with additional training data. Despite this flexibility, this version of our deep network may be currently biased towards common species found along roads, although the algorithms did perform well on static recordings on a range of common and rare species in a range of habitats in the Norfolk Bat Survey (Newson, Evans & Gillings 2015). Nevertheless, in future, extending the training dataset to include annotated bat calls from verified species-call databases to increase geographic and taxonomic coverage, will further improve the generality of our detection tool. Other improvements to the CNN detectors could also be made to lessen taxonomic bias. For example, some bat species have search phase calls longer than the fixed input time window of 23ms of both CNNs (e.g. horseshoe bats). This may limit our ability currently to detect species with these types of calls. One future approach would be to resize the input window (Girshick *et al.* 2014), thus discarding some temporal information, or to use some form of recurrent neural network such as a Long Short-Term Memory (LSTM) (Hochreiter & Schmidhuber 1997) that can take a variable length sequence as input. There are many more unused annotations in the Bat Detective dataset that could potentially increase our training set size. However, we found some variability in the quality of the citizen science user annotations, as in other studies (Kosmala *et al.* 2016). To make best use of these annotations, we need user models for understanding which annotations and users are reliable (Welinder *et al.* 2010; Swanson *et al.* 2016). The Bat Detective dataset also includes annotations of particular acoustic behaviours (feeding buzzes and social calls), which in future can be used to train detection algorithms for different acoustic behaviours (e.g., Prat, Taub & Yovel 2016).

During our evaluation on large-scale ecological monitoring data from Jersey (Walters, Browning & Jones 2016), we demonstrated that our open-source BatDetect CNN_FAST_ pipeline performs as well or better (controlling for spatial and temporal non-independence) compared with an existing widely-used commercial system (SonoBat) that had been manually filtered. Interestingly, although the CNN_FAST_ consistently detected more of the faint and partially-masked calls, most bat passes are likely to still contain at least one call that is clearly-recorded enough to be detected by SonoBat, meaning that the total number of detected bat passes is similar across the two methods. Additionally, our system achieves a massive reduction in the time involved in audio processing - several minutes compared to several days of person-hours (Walters, Browning & Jones 2016). This increase in performance both in terms of speed and accuracy is crucial for future large scale monitoring programmes. Further improvements to our system may come from a better understanding of the patterns of search-phase calls within sequences (Kershenbaum *et al.* 2016). In our example, we used a very simple heuristic to merge individual bat calls into bat passes, but ideally we would also be able to learn this from labelled training data.

## Conclusion

Our BatDetect search-phase bat call detector significantly outperforms existing methods for localising the position of bat search-phase calls, particularly in noisy audio data. It could be combined with automatic bat species classification tools to scale up the monitoring of bat populations over large geographic regions. In addition to making our system available open source, we also provide three expertly annotated test sets that can be used to benchmark future detection algorithms.

## Acknowledgements

We are enormously grateful for the efforts and enthusiasm of the amazing iBats and Bat Detective volunteers, particularly bretarn for the many hours spent providing valuable annotations. We would also like to thank Ian Agranat and Joe Szewczak for useful discussions and access to their systems. This work was supported financially through the Darwin Initiative (Awards 15003, 161333, EIDPR075), Zooniverse, the People’s trust for Endangered Species, Mammals Trust UK the Leverhulme Trust (Philip Leverhulme Prize for KEJ), NERC (NE/P016677/1), and EPSRC (EP/K015664/1 and EP/K503745/1).

## Data Accessibility

All training and test data, including user and expert annotations, along with the code to train and evaluate our detection algorithms will be made available on a Github repository if accepted for publication.

## Supplementary Information

**Supplementary Methods**

**Figure S1** - CNN_FAST_ network architecture description.

**Figure S2 -** Spectrogram annotation interface from Bat Detective (<u>www.batdetective.org</u>).

**Video S1 –** Overview of our system, Bat Detective annotation steps, and sample results.

**Figure S3** – Example search-phase bat echolocation calls from iBats Romania & Bulgaria training dataset.

**Table S1** - Description of BatDetect CNNs test datasets

